# Spatiotemporal multi-omics atlas of the aging neuromuscular junction

**DOI:** 10.1101/2025.11.27.690955

**Authors:** Stefano Amoretti, Samuele Negro, Fabio Lauria, Toma Tebaldi, Giulia Zanetti, Marco Pirazzini, Matthias Mann, Marta Murgia, Gabriella Viero, Michela Rigoni

**Affiliations:** Dept. of Biomedical Sciences, University of Padua, 35131 Padua, Italy; Institute of Biophysics (IBF), CNR Unit at Trento, 38123 Povo, Italy; Laboratory of RNA and Disease Data Science, Department of Cellular, Computational and Integrative Biology (CIBIO), University of Trento, 38123 Povo, Italy; Section of Hematology, Department of Internal Medicine, Yale Comprehensive Cancer Center, Yale University School of Medicine, 06511 New Haven, CT, USA; Myology Center (CIR-Myo), University of Padua, 35131 Padua, Italy; Department of Proteomics and Signal Transduction, Max Planck Institute of Biochemistry, 82152 Martinsried, Germany; Proteomics Program, Novo Nordisk Foundation Center for Protein Research, Faculty of Health and Medical Sciences, University of Copenhagen, 1017 Copenhagen, Denmark

**Author notes:** These authors contributed equally (Stefano Amoretti, Samuele Negro).

**Keywords:** *Neuromuscular junction*, *aging*, *multi-omics*, *post-transcriptional regulation*

## Abstract

The demographic shift toward a global aging population, coupled with rising prevalence of neurodegenerative diseases, poses major public health challenges. Aging is the primary risk factor for most neurodegenerative conditions, making the elucidation of its molecular mechanisms critical for developing effective interventions. Dysfunction of the neuromuscular junction (NMJ), the specialized synapse essential for motor function, is an early hallmark of aging and several neurodegenerative disorders, yet its molecular determinants remain incompletely understood.

To shed light on potential common underlying mechanisms, we performed a spatiotemporal multi-omics analysis of the murine NMJ during aging, uncovering several genes showing decoupling between transcript and protein trajectories that may drive the related progressive motor decline. NMJs of soleus (SOL) and extensor digitorum longus (EDL) muscles, differing in fiber composition and vulnerability to aging and disease, displayed distinct spatiotemporal dynamics: fast-twitch EDL, more susceptible to degeneration, exhibited stronger mRNA-protein decoupling than aged slow-twitch SOL or younger SOL and EDL muscles. Elevated expression of miRNAs and RNA-binding proteins in aged EDL highlights the key role of post-transcriptional regulation in NMJ aging.

This study provides new insights into the physiopathology of neuromuscular aging and offers a resource for investigating mechanisms shared between aging and neurodegenerative diseases. Additionally, it opens avenues for AI-driven discovery of drug targets and early biomarkers, potentially accelerating the development of therapeutic strategies.

## 1 Introduction

By 2050, over 200 million individuals worldwide are expected to suffer from age-related muscular atrophy and weakness, leading to severe socioeconomic and clinical burden (Cruz-Jentoft et al., 2010; Dhillon & Hasni, 2017; Janssen et al., 2004). A hallmark of this geriatric syndrome is the structural degeneration of the neuromuscular junction (NMJ), a process closely linked to muscle weakening and impaired motor function (Carnio et al., 2014; Ham & Rüegg, 2018; Rudolf et al., 2014).

The NMJ is a highly specialized site of nerve-muscle contact, controlling vital functions such as locomotion and breathing. At this junction a motor axon branches, making contacts with multiple muscle fibers. At the contact site, the motor nerve terminal forms varicosities or boutons, from which neurotransmitters are released in the synaptic cleft, triggering muscle contraction. The NMJ undergoes constant remodeling throughout life. In young synapses, a balanced cycle of denervation and reinnervation occurs, maintaining both structure and function of the NMJ. However, with aging, as well as in age-related neurodegenerative disorders, this balance deteriorates, with reinnervation and remodeling becoming eventually overwhelmed by degeneration. Consequently, motor function becomes progressively impaired, leading to motor deficits and muscle atrophy (Jang & Van Remmen, 2011; Li et al., 2011; Taetzsch & Valdez, 2018; Willadt et al., 2018).

A key regulator of longevity across a variety of organisms - from mice to humans - is RNA post-transcriptional regulation (Llewellyn et al., 2024; Tharakan et al., 2021). During aging, regulation of transcripts undergoing translation likely leads to decoupling of mRNA and protein variations, a process that is attracting attention but remains poorly understood (Llewellyn et al., 2024; Solyga et al., 2024). In this context, recent advancements in deep learning, such as the RiboNN model, demonstrate a state-of-the-art capability to predict translation efficiency directly from mRNA sequences, offering a powerful computational avenue to investigate the complex post-transcriptional mechanisms driving mRNA-protein uncoupling in aging and neurodegenerative disorders (Zheng et al., 2024).

To identify potential post-transcriptional controls contributing to the decline in motor performance as the NMJ ages, and which likely come into play also in age-related neurodegenerative diseases, we conducted an in-depth quantitative analysis of age-related transcriptome and proteome dynamics. Using laser capture microdissection (LCM) in predominantly fast (extensor digitorum longus-EDL) or slow-twitch (soleus-SOL) muscles, we collected and pooled NMJs at young (3 months), young adult (12 months) and aged (28 months) stages. We chose these muscles as the NMJs vary in their susceptibility to age-related structural modifications, with the motor unit type being a critical factor. Fast-twitch muscle NMJs, such as those of the EDL, are indeed more susceptible to aging and neurodegeneration compared to slow-twitch counterparts like the SOL (Atkin et al., 2005; Frey et al.,

2000; Hegedus et al., 2007; Murgia et al., 2017; Nijssen et al., 2017). By combining transcriptomics and proteomics, and performing multi-layered analysis, we found robust decoupling of RNA and protein variations as the NMJ ages. The decoupling is particularly relevant in the aged EDL, suggesting the existence of post-transcriptional regulatory mechanisms that may at least partially explain the differences in susceptibilities between the different muscle types.

By providing an atlas with transcriptomes and proteomes of entire NMJs, the present study aims to inform about the neuromuscular synapse as an integrated system, providing a resource to study the effects of aging on NMJ physiology and in age-related neurodegenerative pathologies. The obtained results lay the groundwork for future research on the mechanisms of age-driven decline of the neuromuscular function, and may help AI-driven tools for drug discovery to cope with aging-related diseases of an increasingly older population.

## 2 Methods

### 2.1 Ethical statement

Animals were housed under controlled photoperiod and temperature, with free access to standard food and water. Mice were handled by specialized personnel under the control of inspectors from the Veterinary Service of the Local Sanitary Service (ASL 16-Padua), who are the local officers of the Ministry of Health. The use of animals and the experimental protocols were approved by the Ethical Committee and by the Animal Welfare Coordinator of the “Organismo Preposto al Benessere Animale” of the University of Padua and by the Italian Ministry of Health, and were conducted in accordance with national laws and policies (D.L. n. 26, March 14, 2014), following the guidelines established by the European Community Council Directive (2010/63/EU) for the care and use of animals for scientific purposes.

### 2.2 NMJ transcriptomic profiling

#### 2.2.1 Sample preparation

To identify the NMJs for LCM, C57 male mice weighing approximately 25-30 g at different ages (3, 12 and 28 months old, four mice per time point) were locally injected in the hind limb with 15 µl fluorescent α-BTx-555 diluited in saline (Life Technologies, cat. B35451, 1:200) 1 h before muscle collection. Mice were euthanized with an anesthetic overdose using a mixture of xylazine (48 mg/kg) and zoletil (16 mg/kg) via intraperitoneal injection, followed by cervical dislocation. Next, SOL and EDL were quickly harvested, embedded in OCT, and flash-frozen in liquid nitrogen-cooled isopentane. Cryosections (7 µm thick) were placed onto UV-treated microscope slides. Microdissection was conducted using an automated PALM RoboMover laser microdissector (Carl Zeiss, Oberkochen, Germany). Slides were briefly soaked in 70% ethanol for 30 s to remove the excess of OCT, which could interfere with the analysis. For RNA-seq analysis, 100 NMJs per sample were collected and pooled by LCM within 30 min to preserve RNA integrity. The same procedure was applied to sample collection for NMJ proteomics, except for the ethanol immersion step, which was omitted to avoid potential removal of membrane components. One hundred NMJs/time point were collected by LCM over a 1-hour period.

#### 2.2.2 RNA extraction and sequencing

Total RNA was isolated by incubating LCM samples overnight at 55°C with 50 µl of lysis buffer PKD (Qiagen) and 10 µl of proteinase K (Promega), with tubes inverted. The following day, samples were centrifuged at 10,000 rpm for 10 min, and RNA extracted using the Maxwell® 16 LEV RNA FFPE Purification Kit on the automated Maxwell 16 system (Promega), following the manufacturer’s protocol starting from the DNase treatment step.

For library preparation, cDNA synthesis was performed using the SMARTer Universal Low Input RNA Kit (Clontech Laboratories), following the manufacturer’s instructions. RNA-Seq libraries were prepared with the Nextera XT kit (Illumina), according to the manufacturer’s guidelines. Sequencing was carried out on the Illumina NextSeq 500 platform, pooling up to six libraries per High Output cartridge (300 cycles).

#### 2.2.3 Processing of RNA-Seq data

Nextera adapters were removed (forward: CTGTCTCTTATACACATCTCCGAGCCCACGAGAC, reverse: CTGTCTCTTATACACATCTGACGCTGCCGACGA), the first 11 nucleotides (Cutadapt v.3.7) were trimmed and the reads shorter than 20 nucleotides were discarded. The remaining reads were mapped onto the mouse genome (GRCm39) with STAR (v2.7.10a), using the Gencode M32 gene annotation, based on ENSEMBL release 109. Duplicated reads were removed with Picard Tools MarkDuplicates (v2.26.11) and counts of mapped reads per gene were retrieved by htseq-count --stranded=no from HTSeq (v1.99.2). All programs were used with default settings unless otherwise specified.

Genes not covered by any reads in all samples were removed. To mitigate size differences between libraries, gene expression levels were normalized using the trimmed mean of M-values normalization method implemented in the edgeR Bioconductor package. The normalized count data were modeled using a negative binomial distribution (glmQLFTest function in edgeR), to test for differential expression. The following parameters were considered to define down- and up-regulated genes: absolute value of log2FC < 0.75; statistical significance p-value < 0.05.

#### 2.2.4 Identification of coupled and decoupled genes

Transcriptomic (n = 31,057 genes) and proteomic (n = 903 proteins) datasets were first merged based on gene identifiers, yielding 878 genes common to both datasets. Coupling between transcript and protein levels was determined based on the directionality of fold changes. Briefly, for each gene, the comparison between 3-, 12- and 28-months samples in SOL and EDL was classified according to the concordance between transcriptomic (RNA-seq) and proteomic (mass spectrometry) log2 fold changes. Genes were classified as “coupled” if they exhibited consistent changes (++, --, or ==) and as “decoupled” if changes were discordant (=+, +=, -=, =-, +-, -+) across RNA and protein levels. Only genes with non-missing classifications in both time points were retained for subsequent analysis. This analysis identified 584 genes exhibiting decoupled expression patterns across the different conditions.

#### 2.2.5 Sequence feature analysis

For each gene, GC content and transcript feature lengths - coding sequence (CDS), 3’ untranslated region (3′UTR), and 5’ untranslated region (5′UTR) - were retrieved from Ensembl (GRCm39, release vM32). GC content was computed from the coding sequences of all annotated transcripts using Biostrings v2.70.1 (Bioconductor). CDS and UTR lengths were obtained from annotated transcript models and collapsed at the gene level by selecting the principal transcript. Gene identifiers were matched to coupling classes using the classification described above. To reduce the impact of extreme values, outliers were removed within each muscle, timepoint, and coupling class using the 1.5×IQR rule. Comparisons of GC content, CDS, 3′UTR, and 5′UTR lengths between coupled and decoupled genes were performed separately for each muscle and timepoint using the Wilcoxon rank-sum test, followed by Benjamini–Hochberg (BH) multiple-testing correction. Adjusted p-values were reported as: p < 0.05 (*), p < 0.01 (**), and p < 0.001 (***). Violin plots with overlaid boxplots were generated in R (v4.4) using ggplot2 (v3.5). Outliers were omitted in the visualization for clarity.

#### 2.2.6 Functional enrichment analysis

Annotation enrichment analysis with Gene Ontology terms, REACTOME and KEGG pathways were performed with the ClusterProfiler Bioconductor package. Functional enrichment results from Gene Ontology (GO) analyses were combined across experimental comparisons. A balanced subset of 30 GO terms was selected for heatmap visualization: five top-ranked terms (lowest adjusted p-value) were chosen for each of three main functional categories (*Muscle*, *Neural*, *Extracellular Matrix*) based on keyword-based annotation of GO term descriptions, while the remaining terms were selected from other categories according to significance ranking. Adjusted p-values were converted to enrichment scores as -log₁₀ (p.adjust), clipped at a maximum value of 5 to limit the dynamic range. The resulting matrix of GO terms (rows) and comparisons (columns) was visualized using the ComplexHeatmap package in R. Rows were clustered hierarchically and annotated by functional category. A diverging color scale (blue-white-red) was applied to represent enrichment strength.

GO enrichment of TDP-43 targets was performed using clusterProfiler (v4.10.0) and the org.Mm.eg.db annotation package. Enrichment was tested for Cellular Component ontology, applying a Benjamini-Hochberg adjusted p-value cutoff of 0.01. Enriched terms were visualized with dotplot() (clusterProfiler), where point size represented the number of genes associated with each term and color indicated the adjusted p-value.

#### 2.2.7 Enrichment analysis of regulatory regions

Regulatory enrichment analysis was performed using AURA 2.0 (Dassi et al., 2014), which provides enrichment of RNA-binding proteins and other post-transcriptional regulators over predefined gene sets. Input gene sets included common genes (n=878), decoupled genes (n=584), and genes classified as coupled or decoupled in SOL and EDL comparisons (12m vs 3m, 28m vs 3m). Only regulatory factors enriched (FDR <0.05) in all comparisons were retained for visualization. Bubble plot of regulatory enrichment were generated in R using ggplot2, with dot size proportional to the number of regulated genes and color indicating the FDR-adjusted p-value.

#### 2.2.8 Definition of TDP-43 targets

TDP-43-regulated genes were extracted from the AURA enrichment output, and genes under TDP-43 control were identified for each comparison. Scatter plots displaying the relationship between transcriptomic and proteomic log2 fold changes for TDP-43 targets were generated using ggplot2 and ggrepel, with color-coded uncoupling classes retrieved from the coupling classification described above. Axes were scaled to the same range across all comparisons to allow direct visual comparison. The list of TDP-43 targets significantly enriched at 28 months in EDL was extracted and annotated with gene descriptions from Ensembl (release vM32).

### 2.3 NMJ proteomic profiling

#### 2.3.1 Sample preparation

For NMJ samples, the collection cap of each LCM tube was washed with 10 µl of reduction and alkylation buffer LYSE (PreOmics), the lysate was centrifuged and transferred to 1.5 ml Eppendorf tubes. Samples were further boiled for 5 min to denature proteins, cooled down and sonicated for 10 cycles (1 minute, 50% duty cycle) in a Diagenode Bioruptor. Samples were digested overnight with Lys-C and trypsin, 1 µg/sample at 37°C, under continuous stirring. Peptides were washed using an in house made Stage-Tip centrifuge at 2000 x g, eluted with 60 µl of elution buffer (80% acetonitrile/1% ammonia) into autosampler vials and dried using a SpeedVac centrifuge (Eppendorf, Concentrator plus). Peptides were resuspended in 2% acetonitrile/0.1% TFA before peptide concentration estimation using Nanodrop.

#### 2.3.2 Liquid Chromatography Tandem Mass Spectrometry (LC-MS/MS) analysis

Nanoflow LC-MS/MS analysis of tryptic peptides was conducted on a Q Exactive HF Orbitrap coupled to an EASYnLC 1200 ultra-high-pressure system via a nano-electrospray ion source (all from Thermo Fisher Scientific). Peptides were loaded on a 50 cm HPLC-column (75 μm inner diameter; in-house packed using ReproSil-Pur C18-AQ 1.9 µm silica beads; Dr. Maisch). Peptides were separated using a linear gradient where solvent A was 0.1% formic acid in water and solvent B was 80% acetonitrile and 0.1% formic acid in water, raising the buffer B concentration from 2% to 60% in 100 min, followed by 10 min at 95% B and 10 min at 2% B. The total duration of the run was 120 min. Column temperature was kept at 50°C by a Peltier element-containing, in-house developed oven.

The mass spectrometer was operated in “top-15” data-dependent mode, collecting MS spectra in the Orbitrap mass analyzer (60000 resolution, 300-1650 m/z range) with an automatic gain control (AGC) target of 3E6 and a maximum ion injection time of 20 ms. The most intense ions from the full scan were isolated with an isolation width of 1.5 m/z. Following higher-energy collisional dissociation (HCD), MS/MS spectra were collected in the Orbitrap (15 000 resolution) with an AGC target of 1E5 and a maximum ion injection time of 60 ms. Precursor dynamic exclusion was enabled with a duration of 30 s.

#### 2.3.3 Processing of proteomics data

The MaxQuant software (version 1.6.10.43) was used for the analysis of raw files. Peak lists were searched using the Andromeda search engine (Cox et al., 2011) against the mouse UniProt FASTA reference proteomes version of 2016 for proteomic data and with the gencode. vM32 version for alignment with transcriptome data. Peak lists were also searched against a common contaminants database. Carbamidomethyl was included in the search as a fixed modification, oxidation (M) and acetyl (Protein N-term) as variable modifications. The FDR was set to 1% for both peptides (minimum length of 7 amino acids) and proteins and was calculated by searching a reverse database. Peptide identification was performed with an initial allowed precursor mass deviation up to 7 ppm and an allowed fragment mass deviation 20 ppm.

Proteomic data were based on raw abundances calculated from the area under the curve (AUC) for each peptide-to-spectrum match (PSM), derived from mass-to-charge ratio (m/z), retention time, and the number of detection events. Preliminary filtering steps removed peptides with low or inconsistent abundance, defined as those not consistently present across all replicates within at least one condition, as well as peptides identified as contaminants.

The proteomics data were processed using the *ProTN* pipeline (https://github.com/TebaldiLab/ProTN), a comprehensive workflow specifically designed for downstream analysis of label-free DDA proteomics data derived from both Proteome Discoverer (PD) and MaxQuant (MQ) outputs. The pipeline integrates all key post-processing steps, including normalization, imputation, summarization, and differential expression analysis, at both the peptide and protein levels. First, raw intensity values were log2-transformed and normalized separately at peptide and protein levels to account for systematic variability. At the peptide level, median-centering was performed using the equalMedianNormalization function, which adjusts each sample distribution to a common median value. At the protein level, peptide intensities were summarized into protein abundances via the *median sweeping* method (medianSweeping function), which combines median normalization with robust peptide-to-protein aggregation, ensuring accurate quantification despite varying numbers of peptides per protein. Missing values were imputed using the *PhosR* algorithm (Kim et al., 2021) which applies a condition-aware imputation strategy that preserves biological variability while minimizing artefactual biases. As a fallback, Gaussian imputation was applied in rare cases where PhosR could not resolve the missing data structure. For differential expression analysis, ProTN performs statistical testing separately for peptides and proteins. Protein-level differential expression was assessed using *DEqMS* (Zhu et al., 2020), a method that extends the *Limma* framework (Ritchie et al., 2015), by incorporating peptide-level evidence and modeling variance as a function of peptide count per protein, thereby improving accuracy especially for proteins supported by few peptides. Peptide-level testing was performed directly using *Limma*. For each protein or peptide, fold change, p-values, adjusted p-values (FDR), and log2 expression values were calculated. Proteins and peptides were classified as significantly up- or down-regulated only if they passed multiple thresholds for fold change, statistical significance, and signal intensity. The following parameters were considered to define down- and up-regulated proteins: absolute value of log2FC < 0.75; statistical significance p-value < 0.05.

### 2.4 Immunofluorescence

Cryosections of SOL and EDL not designated for omics analysis were stored at - 80°C for histological examinations. Once thawed at RT, cryosections were fixed in 4% PFA in PBS for 30 min at RT, and quenched in 0.24% NH4Cl PBS for 20 min. After permeabilization and 2 h saturation in blocking solution (15% goat serum, 2% BSA, 0.25% gelatine, 0.20% glycine, 0.5% Triton-X100 in PBS), samples were incubated with primary antibody against Ftl1 (Abcam, cat. ab69090, 1:200) and Dcn (Novus biologicals, cat. NBP1-57923, 1:200) for 72 h in blocking solution at 4°C. Following 3 washes with PBS, sections were incubated with secondary antibodies and α-BTx AlexaFluor-555 to stain post-synaptic acetylcholine receptors (AChRs), diluted in PBS +0.5% Triton X-100 for 2 h at room temperature. After 3 additional washes in PBS for 10 min, samples were mounted using Dako fluorescence mounting medium (Agilent Technologies, cat S3023). Images were collected using a Zeiss LSM 900 Confocal microscope equipped with a 40× HCX PL APO NA 1.4 oil immersion objective. Laser excitation, power intensity, and emission ranges were optimized for each fluorophore to minimize cross talk.

## 3 Results

### 3.1 Age-related changes in the transcriptome and proteome of SOL and EDL NMJs

NMJs from young (3 m), young adult (12 m), and old (28 m) mice were isolated from SOL (slow-twitch) and EDL (fast-twitch) muscles by LCM, followed by RNA extraction and sequencing (transcriptome), or protein purification and mass spectrometry profiling (proteome) (Figure 1A). A representative immunostaining of young and old NMJs is reported in Figure 1B, revealing age-dependent structural alterations in the aged neuromuscular synapse, such as NMJ fragmentation and partial denervation (Willadt et al., 2018).

**FIGURE 1.**
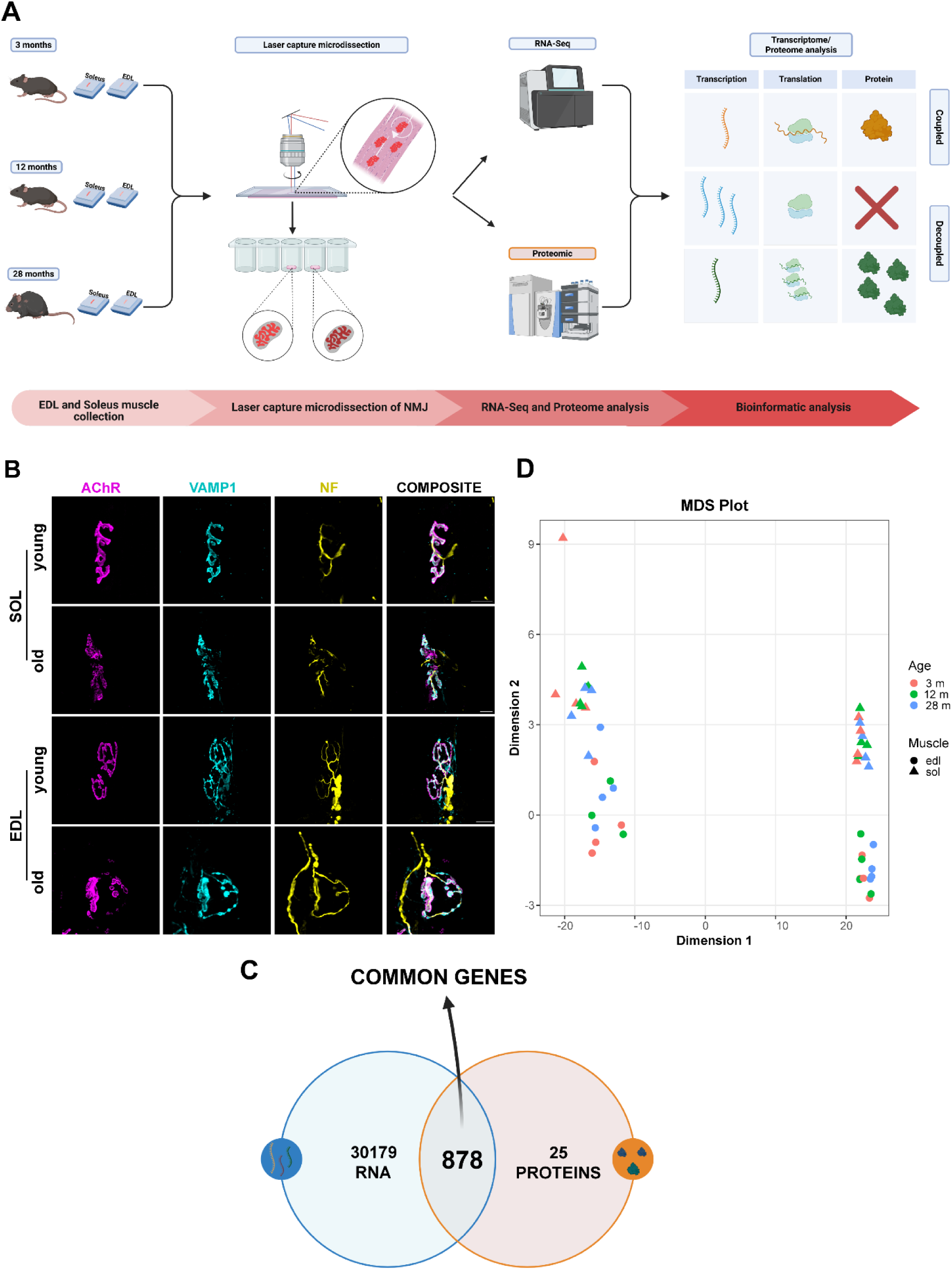
Generating a multi-omics atlas of NMJ aging. (A) Experimental workflow. (B) Representative immunostaining of neuromuscular synapses from young (3 m) and old (28 m) SOL (upper panels) and EDL (lower panels). The post-synaptic compartment is identified by fluorescent α-BTx (magenta), the presynaptic nerve terminal and the axon by antibodies against VAMP1 (cyan) and neurofilament (NF) (yellow), respectively. Scale bars: 20 μm. (C) Overlap between RNAs, common genes and proteins. (D) Multi-omics MDS plot of aging NMJs, showing a clear separation between proteomic and transcriptomic abundances, muscle types and different ages.

A total of 31,057 genes and 903 proteins were identified. To reveal muscle or age specific similarities across groups, we employed multidimensional scaling (MDS), which shows that SOL and EDL samples generate distinct clusters in both the transcriptome (Figure S1A) and proteome (Figure S1B). Precisely, inter-muscular protein levels show clear separation of muscles at all time points (Figure S1B), whilst transcriptional levels are less pronounced (Figure S1A).

The intersection of transcriptomics and proteomics data reveals 878 overlapping genes (Figure 1C). The MDS plot of these common genes confirms the distinct clustering of EDL and SOL samples at both levels and highlights a comprehensive difference in RNA vs protein abundances (Figure 1D). These findings indicate that, overall, muscle-specific differences in gene expression outweigh age-specific changes, and that proteomes provide a more accurate representation of muscle specificity than transcriptomes.

To identify the inter-muscular differences within the same age (e.g. SOL 3 m vs EDL 3 m), and the intra-muscular differences over time (e.g. SOL 3 m vs SOL 28 m), we calculated the differentially expressed genes (DEGs) and proteins (DEPs). The analysis confirms the above-mentioned inter-muscular differences, underscoring the unique transcriptome of the aged EDL. Indeed, at 28 months, substantial transcriptional changes occur in the EDL, displaying hundreds of significantly upregulated and downregulated genes (Figure S1C). Despite persistent inter-muscular differences at the transcriptional level - resembling those observed in humans (Abbassi-Daloii et al., 2023) - these alterations not only endure over time but also intensify at the protein level (Figure S1D).

### 3.2 Protein-RNA decoupling increases during aging

Our multi-omics atlas has the unique potential to uncover muscle-specific post-transcriptional mechanisms regulating gene expression during aging, which may alter the direct proportion between transcript and protein abundance (Schwanhäusser et al., 2011; Tebaldi et al., 2012). By directly comparing mRNA vs protein abundance on a genome-wide scale (see Methods), we tested the hypothesis that post-transcriptional regulation contributes to the muscle-specific differences revealed at the proteome level.

We identified a subset of 584 genes with significantly non-concordant transcript and protein changes, referred to as ‘decoupled genes’ (Figure 2A). The representative scatter plot in Figure 2B compares the fold changes of transcriptional and protein profiling, where grey dots represent genes with coupled expression and the others those that are decoupled. The latter were further categorized into different classes and color-coded according to the type of change (e.g. upregulated at the transcript level but downregulated at the protein one). In the EDL, the number of decoupled genes increases with age, reaching nearly four times the level observed in the SOL at 28 months (192 vs 53) (Figure 2C, middle and Figure 2E). Conversely, inter-muscle differences remain comparatively stable across ages (Fig. 2C, right). This finding highlights a stronger intra-muscle age-dependent shift toward decoupling in the EDL.

**FIGURE 2.**
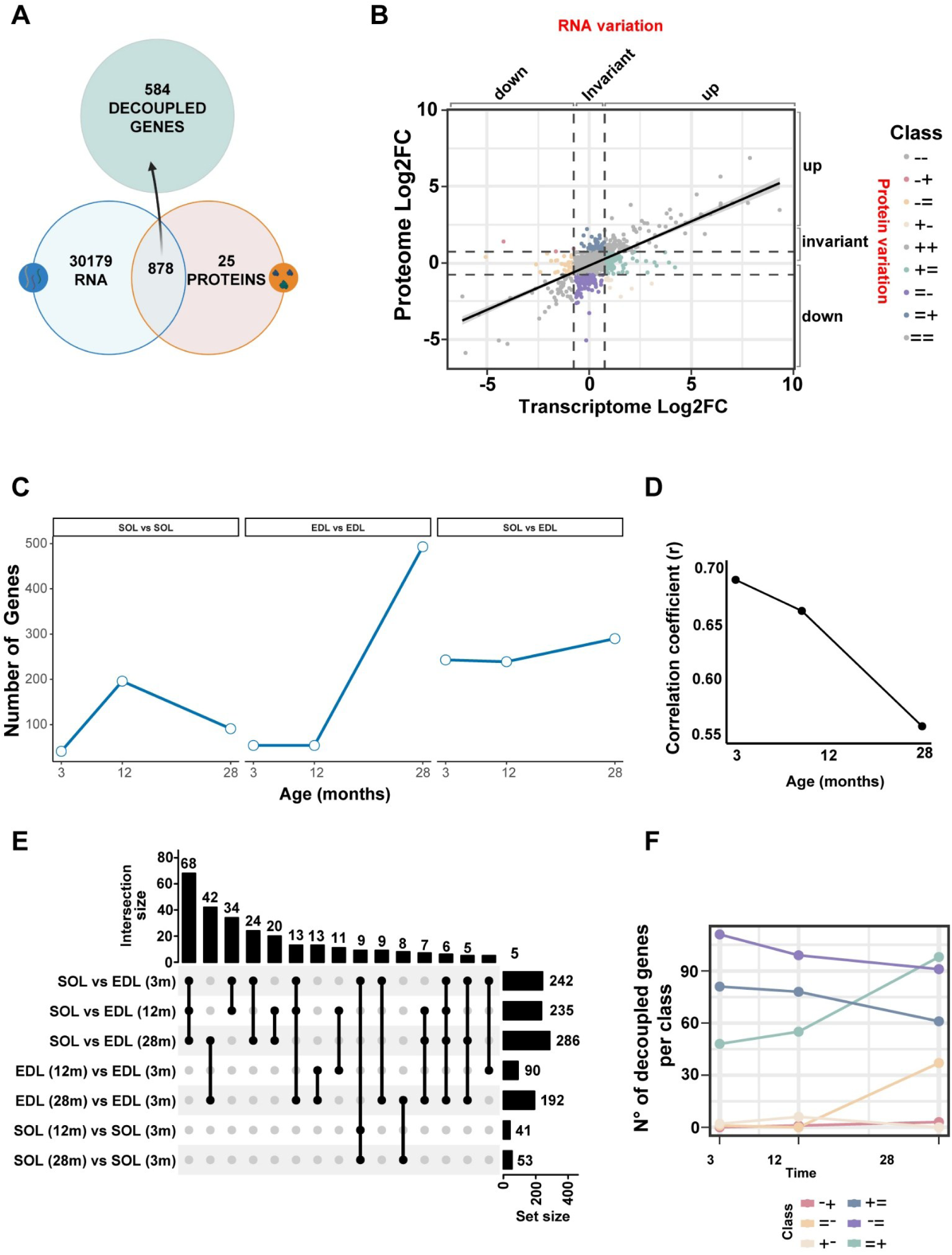
Decoupling in gene expression increases during aging. (A) Venn diagram showing the overlap between transcripts and proteins across all samples. 584 genes display discordant RNA-protein behavior across muscles and ages, and are referred to as ‘decoupled’. (B) Scatter plot comparing log2 fold changes (Log2FC) in transcriptome and proteome for a representative condition, illustrating distinct classes of RNA-protein relationships (color-coded by direction of change at RNA and protein level). The black line indicates the global correlation between the two datasets. (C) Line plot showing the number of decoupled genes in SOL and EDL across ages. Decoupling progressively increases with age, particularly in the EDL. (D) Correlation coefficients between RNA and protein log2FC values in the two muscles across ages, showing a gradual decline in RNA-protein coupling from 3 to 28 months. (E) UpSet plot displaying intersections of decoupled gene sets across muscles and time points. Notably, the degree of decoupling at 28 months is nearly fourfold higher in the EDL than in the SOL (192 vs 53 genes), and about one-third of decoupled genes (182) are shared between the two muscles. (F) Line plot showing the distribution of decoupled genes among the different RNA-protein classes (color-coded as in panel B).

The increased decoupling during aging is further supported by the declining correlation between the changes in transcriptome and proteome levels in the inter-muscle difference (Figure 2D), which decreases from r=0.69 at 3 months (Figure S2A), to r=0.662 at 12 months (Figure S2B), and r=0.558 at 28 months (Figure S2C). A consistent pattern of discordant expression was observed across all age groups, with significant shifts in decoupling categories as the NMJ ages (Figure S2D-F). Specifically, when comparing decoupled genes at the protein and transcript levels (classes =+ and =-), we found that across all time points most protein-level changes could not be explained by corresponding mRNA alterations (purple and cadet blue lines in Figure 2F). In aged muscles, however, there is an enrichment in genes displaying transcriptional changes without parallel alterations in protein abundance (classes += and -=, green and light orange lines in Figure 2F, respectively), suggesting that changes in transcriptional levels in both directions (up/down) intensifies with aging, whereas the proteome remains comparatively stable.

We validated key findings by immunofluorescence. Two candidate genes displaying highly decoupled expression patterns and likely contributing to adaptive processes at the aged NMJ (Brandan & Gutierrez, 2013; Halon-Golabek et al., 2019; Mietto et al., 2021; Ryan et al., 2020) were selected: Decorin (Dcn) and Ferritin light chain (Ftl1). These genes were identified as decoupled because their transcripts remain stable while their corresponding protein levels exhibit a marked upregulation in the aging NMJ (Figure 3 panels A,C). Representative immunostaining of Dcn and Ftl1 confirms the increased protein signal over time at both the SOL (left) and the EDL (right) NMJs (panels B and D, respectively). These results support the conclusion that significant regulatory mechanisms operate beyond transcription to shape the NMJ proteome during aging.

**FIGURE 3.**
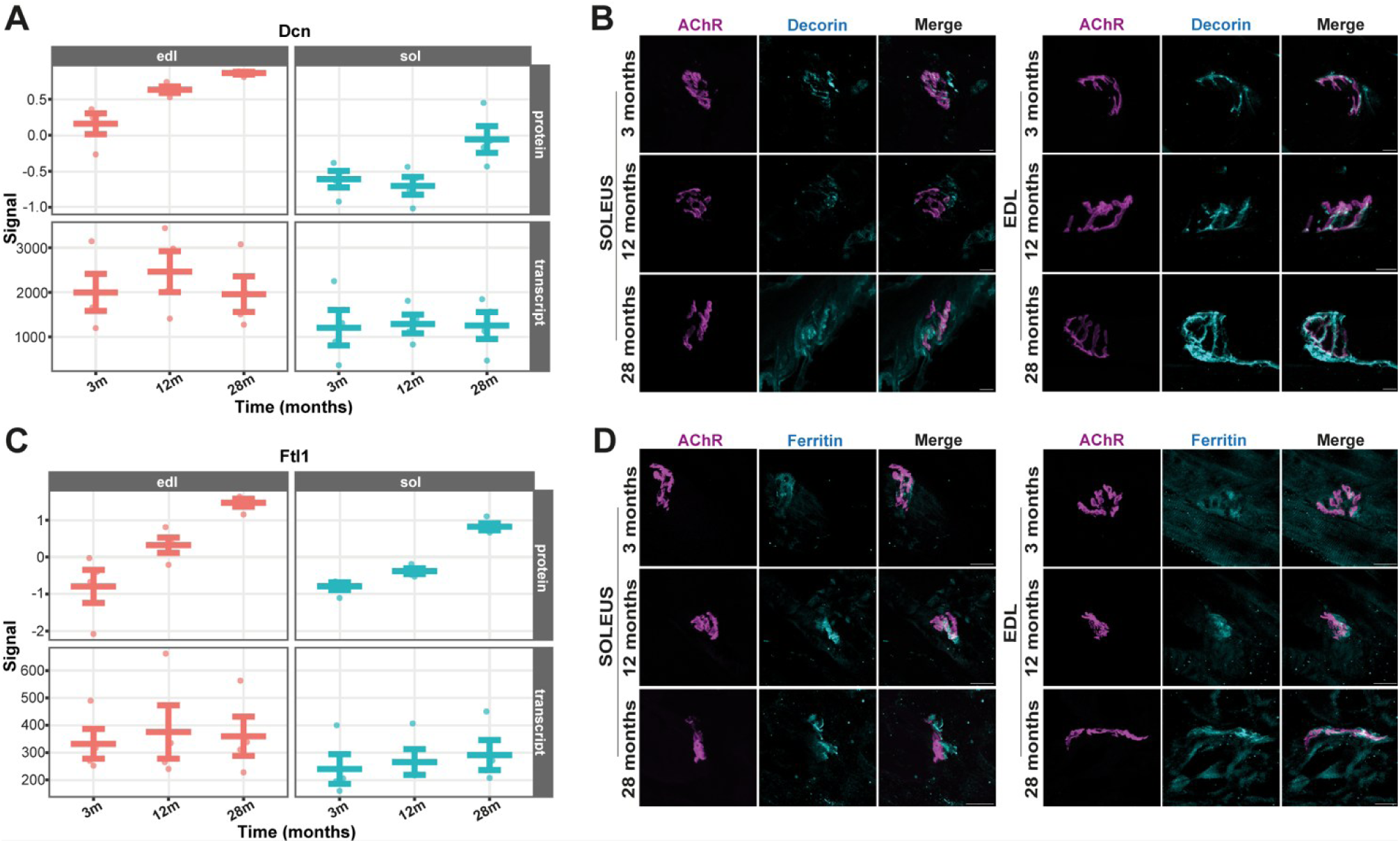
Validation of decoupling and specific increase of protein expression of Decorin (Dcn) and Ferritin light chain (Ftl1) at aged NMJs. (A) Dcn expression at protein and transcript levels. (B) Dcn protein expression shown by immunofluorescence at SOL (left) and EDL (right) NMJs during aging. Acetylcholine receptors (AChR) are in magenta, Dcn in cyan. (C) Ftl1 expression at protein and transcript levels in the different muscles during aging. (D) Ftl1 protein expression by immunofluorescence at SOL (left) and EDL (right) NMJs during aging. AChR are in magenta, Ftl1 in cyan. Scale bars: 10 μm.

### 3.3 Multi-omics signature of aging reveals early decline of contractile programs in the SOL and late-onset neural stress in the EDL

To characterize the molecular pathways underpinning NMJ aging, we performed Gene Ontology (GO) enrichment analysis of decoupled genes derived from temporally resolved comparisons in SOL and EDL muscles. The top 30 most significantly enriched biological processes (-log10 adjusted p-value) were selected, ensuring balanced representation across muscle, neural and extracellular matrix (ECM) pathways. The GO terms visualized as a heatmap show enriched biological processes in both muscles during aging (Figure S3). The heat map delineates a clear divergence between SOL and EDL aging responses. In the SOL, contractile-related GO terms - including sarcomere, myofibril and muscle contraction - are weakly enriched at earlier stages (e.g., 12 m vs 3 m), but display strong enrichment at a later stage (28 m). This pattern suggests a delayed reinforcement of contractile programs, potentially reflecting late-onset compensatory remodeling or increased mechanical stress in this aging postural muscle. In stark contrast, EDL exhibits a markedly different trajectory, with neural-associated processes - e.g. synapse, axon and post synapse - becoming significantly enriched in the latest aging stage (28 m vs 3 m). This late-onset activation of neuronal pathways may reflect denervation phenomena or neuromuscular destabilization, consistent with the known susceptibility of EDL (Deschenes et al., 2010). ECM-related terms, including collagen-containing extracellular matrix and basement membrane organization, exhibit transient enrichment in intermediate SOL comparisons, suggesting a temporally confined window of matrix remodeling, which is absent or diminished in the EDL.

### 3.4 Post-transcriptional control of gene expression at the NMJ during aging

To further dissect the mechanisms underlying transcript-protein decoupling during aging, we explored whether some of the most well known *cis*-regulatory features of transcripts and *trans*-acting factors, such as RNA Binding Proteins (RBPs) and microRNAs (miRNAs) could influence the observed decoupling.

As a proxy for transcript stability, we examined the GC content and the length of the coding sequence (CDS) and 3′/5′UTRs in both muscles and coupled and decoupled transcripts. At 28 months, decoupled genes displayed significantly higher GC content than coupled genes in the aged SOL muscle (Figure 4A, first row), a feature typically associated with increased mRNA stability and reduced degradation. The CDS length was also consistently longer in decoupled genes, a feature particularly evident at 28 months in both muscles (Figure 4A, second row). In the EDL, the 3′UTR was significantly longer in decoupled genes at 28 months (Figure 4A, third row). This result is in agreement with the fact that longer 3′UTRs increase the number of potential binding sites for RNA-binding proteins (RBPs) and microRNAs (miRNAs), potentially increasing post-transcriptional regulation (Mayr 2019; Mayya & Duchaine, 2019; Zhao et al., 2011). Conversely, no significant difference was observed in the 5′UTR length at either time points in both muscles (Figure 4A, fourth row). Collectively, these findings suggest that sequence features associated with increased mRNA stability and regulatory complexity (such as high GC content and longer CDSs and 3′UTRs) are selectively enriched in decoupled genes during aging. This highlights a specific characteristic of the aging transcriptome, as these genes are likely more susceptible to post-transcriptional regulation. Consequently, the decoupling of protein levels from RNA abundance may occur with higher frequency in aging muscles.

**FIGURE 4.**
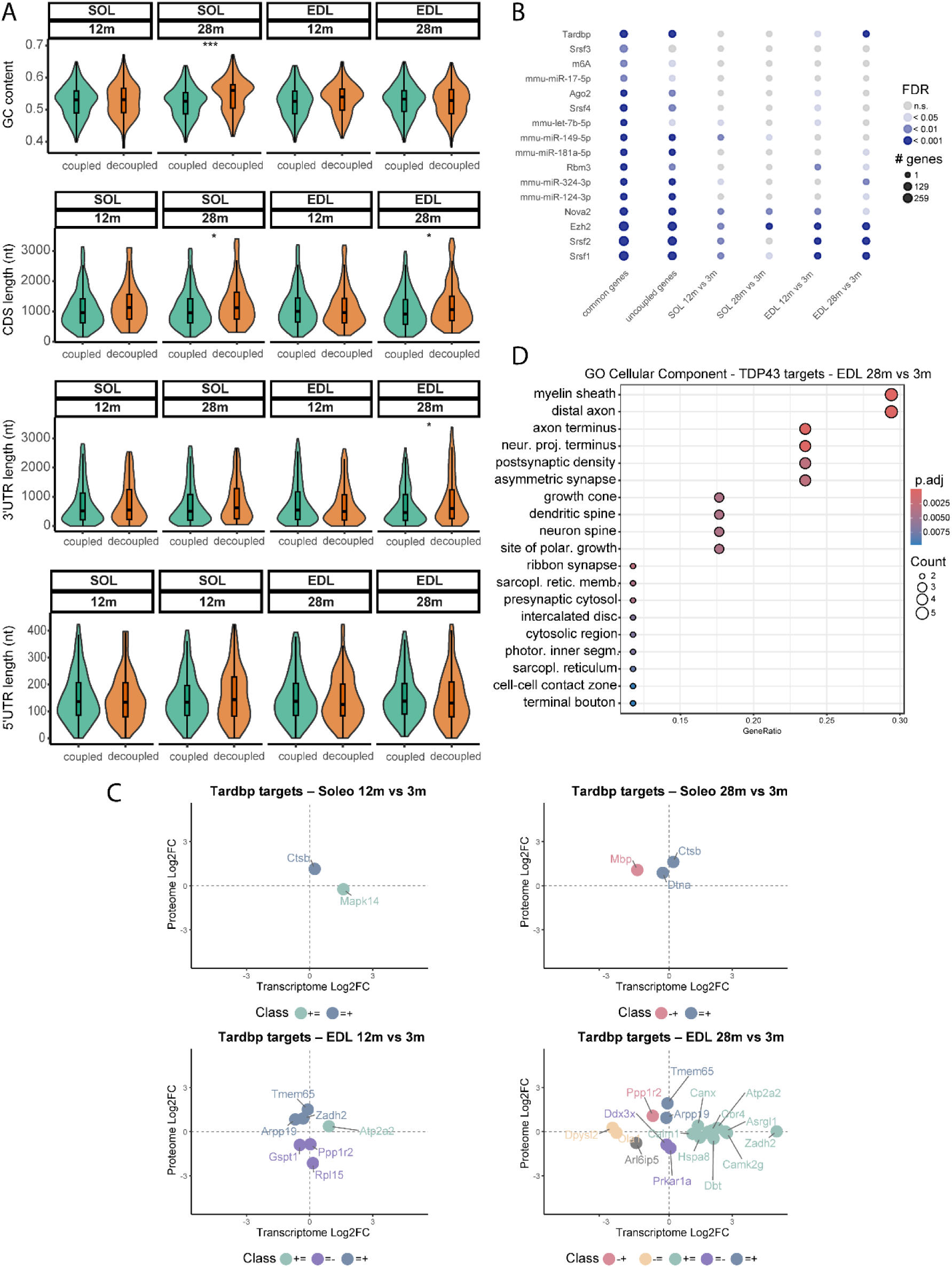
Post-transcriptional features and regulators of transcript-protein uncoupling during aging. (A) Violin plots comparing GC content, CDS length, and 3′/5′UTR length between coupled and decoupled transcripts in SOL and EDL at 12 and 28 months. (B) Regulatory enrichment analysis based on the AURA database. Circle size represents the number of regulated genes, while color indicates FDR; TDP-43 (Tardbp) emerges as the most significantly enriched RBP in aged EDL. (C) Scatter plots of transcriptome (x-axis) versus proteome (y-axis) fold changes for TDP-43 targets across aging comparisons. Genes are color-coded by decoupling class, with several targets in aged EDL (28 m vs 3 m) showing strong RNA-protein decoupling. A subset of these genes has also been implicated in ALS (see Table 1), highlighting their potential relevance for neuromuscular aging. (D) Gene Ontology analysis of TDP-43 targets in aged EDL reveals enrichment for neuronal and mitochondrial components. Dot size represents the number of genes annotated to each term, and the color scale reflects adjusted p-values.

**TABLE 1.**
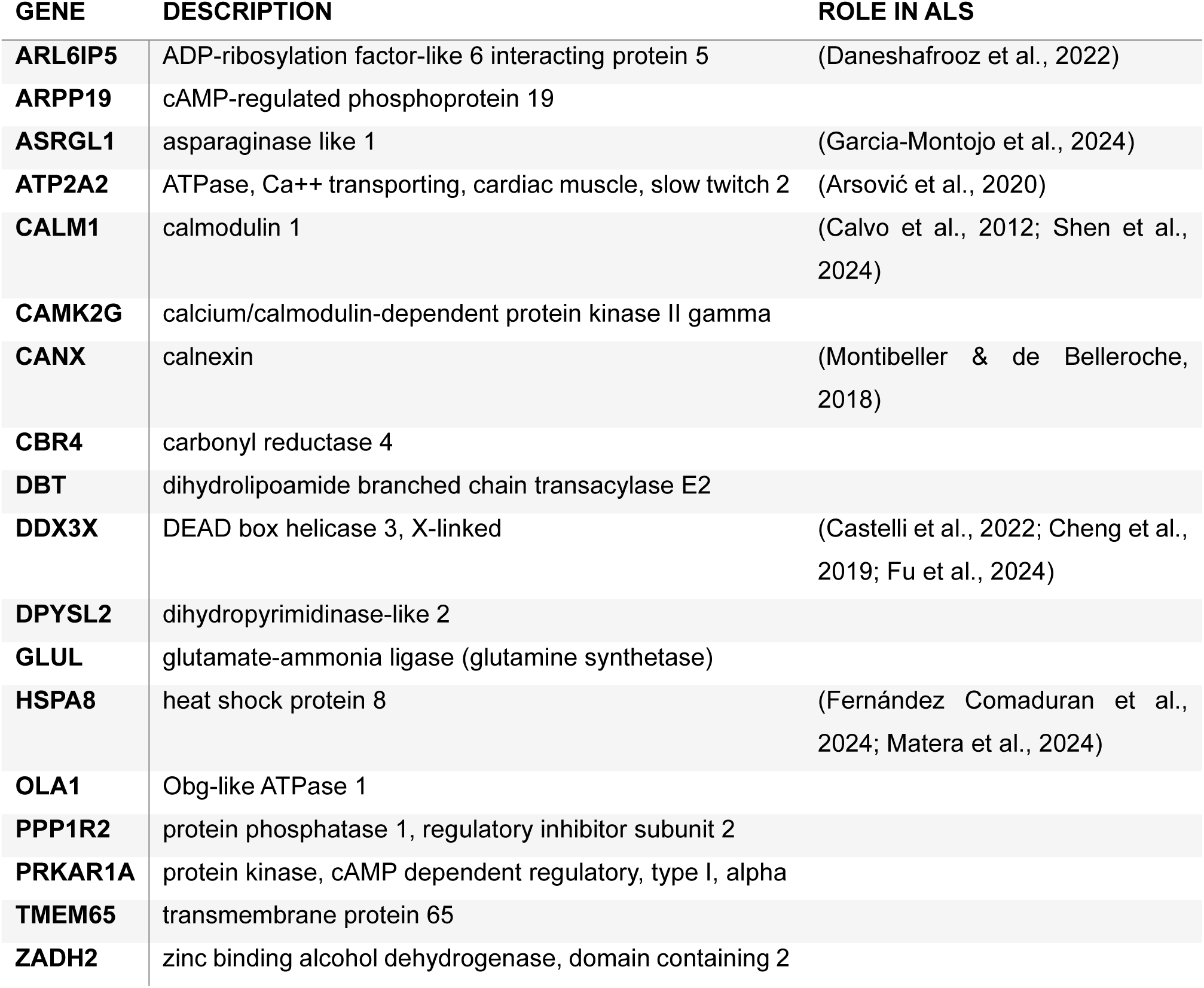
TDP-43-regulated genes in the EDL at 28 months of age. Listed are TDP-43 targets detected in the aged EDL, along with their putative roles in ALS as reported in literature.

Next, we asked whether *trans*-acting factors contribute to driving age-related decoupling and focused on 3′UTR-mediated interactions with RBPs and miRNAs. Querying the AURA database (Dassi et al., 2014) revealed a significant enrichment of known post-transcriptional regulators binding the decoupled transcripts, particularly in aged EDL muscle (Figure 4B). Among the RBPs, the TAR DNA-binding protein 43 (TDP-43) - a key player in Amyotrophic Lateral Sclerosis (ALS), Frontotemporal dementia and hallmark of various neurodegenerative disorders (Buratti, 2001; Kabashi et al., 2008; Kwong et al., 2007; Suk & Rousseaux, 2020; Zeng et al., 2024) - was strongly enriched at 28 months compared to 3 months, suggesting an increased susceptibility of aging muscle to TDP-43-mediated regulation of mRNA homeostasis. Interestingly, other regulators displayed opposite or distinct patterns: for example, Nova2 was more prominent in younger muscle, while Rbm3 and Mbnl2 remained consistently represented across ages. This finding is particularly intriguing, -òà as TDP-43 is known to form aggregates in ALS, a devastating age-related neuromuscular disease (Josephs et al., 2014; Ling et al., 2013; Neumann et al., 2006; Prasad et al., 2019). Notably, the enrichment of TDP-43-mediated interactions occurs in the EDL, the muscle type preferentially affected in ALS, but not in the SOL, which is relatively spared from the disease (Hegedus et al., 2007). This concordance between disease susceptibility and age-related post-transcriptional regulation strengthens the link between aging-associated decoupling and neuromuscular vulnerability.

To further explore the functional impact of TDP-43 binding on age-related decoupling, we examined the subset of transcripts identified as TDP-43 targets by AURA. Scatter plots of transcriptome versus proteome fold changes (Figure 4C) confirm that the extent of the decoupling of TDP43 targets was more pronounced in EDL at 28 months, where multiple TDP-43 targets displayed strong post-transcriptional regulation. The genes are colored by their decoupling class, highlighting distinct patterns of regulation across muscles and ages. Importantly, Table 1 lists the TDP-43-regulated genes identified in aged EDL (Fig. 4C, lower-right plot), including references related to their involvement in ALS. The overlap with ALS-associated genes underscores that these aging-dependent TDP-43 targets constitute biologically relevant candidates, warranting further investigation into their contribution to neuromuscular vulnerability.

Finally, to identify enriched molecular functions of TDP-43 targets in aging muscles, we performed a Gene Ontology enrichment analysis focusing on the EDL at 28 months (Figure 4D). The analysis reveals neuronal and mitochondrial components as the most affected categories. Terms such as “myelin sheath,” “distal axon,” “growth cone,” and “postsynaptic density” point to a robust involvement of axonal transport and synaptic maintenance, while mitochondrial-related terms emphasize oxidative metabolism and membrane dynamics. These findings suggest that TDP-43-driven post-transcriptional decoupling preferentially targets genes essential for neuromuscular junction stability, axonal integrity and mitochondrial homeostasis, hallmarks of age-related denervation and impaired energy metabolism.

Together, these results showcase post-transcriptional decoupling as potentially relevant process during muscle aging arises from both *cis*-encoded sequence features and *trans*-acting regulatory mechanisms. By selectively acting on transcripts with long 3′UTRs and stable GC-rich coding regions, TDP-43 emerges as a key regulator of age-associated dysregulation, possibly exacerbating the functional decline of vulnerable fast-twitch fibers through impaired synaptic maintenance and altered metabolic homeostasis.

## 4 Discussion

The NMJ serves as the critical hub of motor neurons and skeletal muscles, ensuring proper movement and muscle control. As aging progresses, structural and functional changes at the NMJ contribute to reduced synaptic efficiency, impaired neurotransmission, muscle atrophy, and reduced regenerative potential of the neuromuscular system (Hastings et al., 2023; Painter et al., 2014; Willadt et al., 2018). The present study provides a multi-omics atlas that can help identify the post-transcriptional regulatory mechanisms that shape the changes in the NMJ proteome during aging, thereby elucidating the molecular basis of the progressive anatomical and functional decline of the neuromuscular synapse. Beyond advancing our understanding of the biology of normal aging, this study delivers a powerful tool for shedding light on age-related neurodegenerative disorders, where NMJ dysfunction - such as that occurring in ALS - is a key pathological hallmark.

We proved the power of this resource showing inter-muscular differences between neuromuscular synapses from predominantly slow-twitch (SOL) and fast-twitch (EDL) muscles at both transcriptional and, more markedly, proteomic level. In addition, we showcase its use finding unique molecular properties of these muscles, and confirming that muscle-specific differences outweigh age-specific changes (Murgia et al., 2017; Naruse et al., 2023). A key finding is the identification of genes with discordant expression patterns between transcripts and the encoded proteins, referred to as "decoupled". Importantly, decoupling increases with aging, particularly in the EDL, pointing to a tight post-transcriptional regulation coming into play to shape the NMJ proteome during aging, which would have not emerged through transcriptional or proteomics analyses alone. Our functional analyses further delineate two distinct aging strategies in skeletal muscle: SOL engages a delayed, yet muscle-intrinsic transcriptional program primarily centered on contractile structure and transient ECM remodeling, while EDL undergoes a later, neuronal-driven transcriptional shift, consistent with neuromuscular destabilization. These differences highlight the importance of muscle-type specificity in the aging process and suggest divergent vulnerabilities and compensatory capacities across muscle subtypes.

To validate the multi-omics dataset, we conducted a targeted analysis of protein expression for 2 decoupled genes, Dcn and Ftl1. These genes were selected based on: i) their decoupled expression profiles, characterized by stable transcript levels and marked upregulation at the protein level, thereby supporting the presence of regulatory mechanisms beyond transcription, and ii) their likely involvement in adaptive or compensatory responses at the aged NMJ, potentially contributing to the maintenance of synaptic integrity and function despite age-related challenges. Indeed, Dcn helps maintain skeletal muscle integrity and support long-term muscle repair by regulating satellite cell behavior, essential for muscle regeneration, thus contributing to NMJ longevity and function (Brandan & Gutierrez, 2013). Notably, Dcn expression increases with aging, especially in the EDL.

Ferritin, a protein responsible for binding and safely storing iron, also plays crucial roles in skeletal muscle physiology. Iron is essential for energy metabolism in neurons and is required for myelination. In aging and disease models, ferritin and iron storage proteins like Ftl1 and Fth1 are elevated in both skeletal muscle and neurons. Similarly, we report an age-related increase in Ftl1 expression at both EDL and SOL NMJs (Halon-Golabek et al., 2019; Mietto et al., 2021; Ryan et al., 2020), confirming the value of our dataset as a resource with broad applications.

The increased gene-protein decoupling with age emerged in our study aligns with recent large-scale human proteomic analyses showing widespread transcript-protein decoupling with age across multiple tissues (Ding et al., 2025), supporting the notion that post-transcriptional regulation is a conserved feature of the aging process. As a broad concept, this phenomenon can be interpreted in two contrasting ways: i) as an adaptive response that allows cells to maintain homeostasis under stress by fine-tuning protein output independently of transcription, or ii) as a maladaptive alteration that disrupts the coordination of gene expression and protein synthesis, thereby contributing to the progressive decline in cellular and tissue function (Lai et al., 2024; Solyga et al., 2024). Whether and how age-dependent post-transcriptional changes contribute to disease susceptibility is an open question.

Among trans-acting factors, TDP-43 emerges as the most prominent regulator of decoupled transcripts in the aged EDL. TDP-43 is a highly conserved RBP involved in mRNA stability, splicing and transport, whose dysregulation is implicated in neurodegenerative diseases such as ALS (Kabashi et al., 2008; Kwong et al., 2007; Suk & Rousseaux, 2020). TDP-43 binding is selectively enriched in this muscle type, which is also preferentially affected in ALS (Deschenes et al., 2010), while it is largely absent in the relatively spared SOL. Other RBPs - including Nova2, Rbm3, and Mbnl2 - show age-independent or distinct patterns, highlighting the specificity of TDP-43 involvement. Notably, several TDP-43 targets overlap with ALS-associated genes, strengthening the translational relevance of our findings and suggesting that aging-related post-transcriptional dysregulation may converge with disease mechanisms. Functional annotation of TDP-43 targets indicates a predominant involvement of neuronal and mitochondrial components, encompassing processes such as axonal transport, synaptic stability, and oxidative metabolism (Altman et al., 2021; Briese et al., 2020). These pathways are central to NMJ integrity and energy homeostasis and decline with age. We propose that TDP-43-mediated decoupling contributes to functional deterioration by selectively impairing transcripts essential for maintaining synaptic and metabolic balance, particularly in fast-twitch fibers prone to degeneration.

In summary, this study presents a comprehensive omics data atlas of the NMJ, offering a valuable resource for advancing our understanding of the molecular and cellular processes underlying normal aging. Beyond aging, the atlas provides a framework for exploring mechanisms involved in neurodegenerative disorders in which NMJ deterioration is a hallmark, such as ALS. Together, these findings underscore the utility of integrative, multi-omics approaches in uncovering the complex interplay between structure, function, and disease vulnerability at the NMJ.

## Acknowledgements

This work was supported by AFM Telethon (to MP) and by the University of Padua (to MR and MP).

## Author contributions

MR conceived and supervised the study, with the contribution of MP, GV and MMu. SA performed the bioinformatic analysis under the supervision of FL, TT and GV. MP provided the NMJ transcriptomic dataset. SN and GZ performed LCM for NMJ proteomics. MMu performed the proteomic analysis. MMa provided resources. SN performed data validation. SA, MR, GV wrote the manuscript, with comments and suggestions by all authors.

## Disclosure and competing interests statement

No competing interests

## Data Availability

Raw and analyzed RNA-seq data generated in this study have been deposited in the Gene Expression Omnibus (GEO) under the accession code GSE310629.

The mass spectrometry proteomics data have been deposited in the ProteomeXchange Consortium via the PRIDE partner repository with the dataset identifier PXD056155.

## Notes

### Competing Interest Statement

The authors have declared no competing interest.

