## Supplementary Materials for "Spatiotemporal multi-omics atlas of the aging neuromuscular junction"

SUPPLEMENTARY FIGURE 1

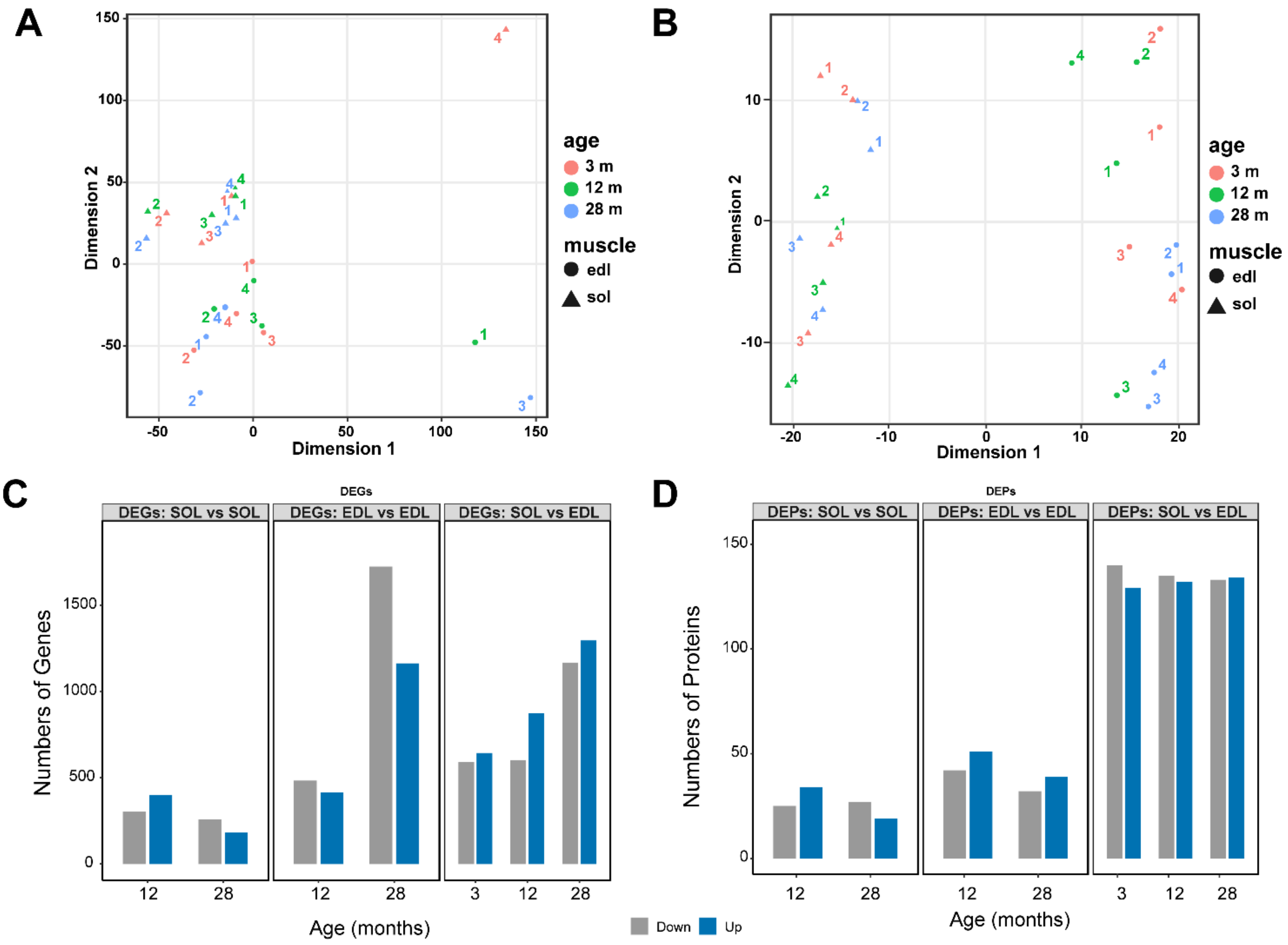


**FIGURE S1**. The NMJ transcriptomes and proteomes of SOL and EDL are different across life span. Differences between SOL and EDL NMJs during aging occur at the transcriptional level (A, C), and are even more pronounced at the proteomic one (B, D). Number of differentially expressed genes (DEGs, panel C) and proteins (DEPs, panel D) (up- and down-regulated), identified based on a fold change (FC) threshold of FC > 0.75 (upregulated) or FC < -0.75 (downregulated), with a statistical significance threshold of p-value < 0.05.

SUPPLEMENTARY FIGURE 2

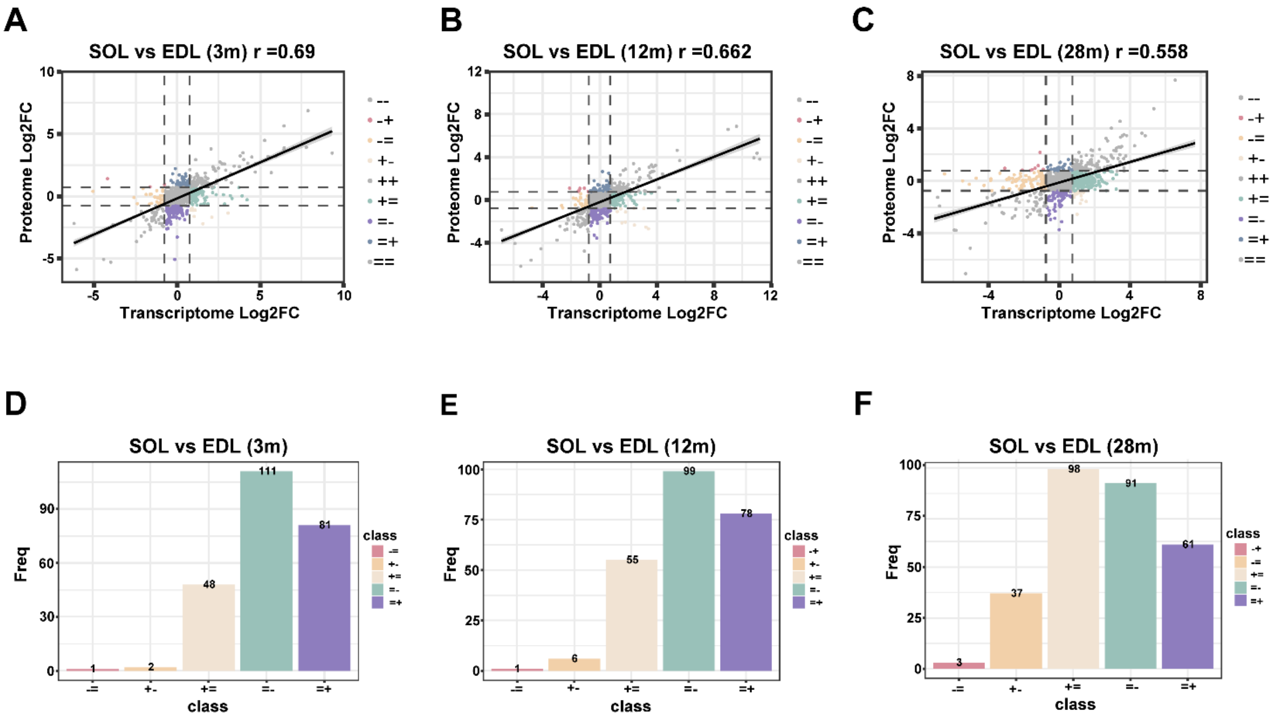


**FIGURE S2**. Scatter plots of decoupled genes in the inter-muscular difference during aging and relative bar plots showing number of decoupled genes per class at 3 months (A, D), 12 months (B, E) and 28 months (C, F).

SUPPLEMENTARY FIGURE 3

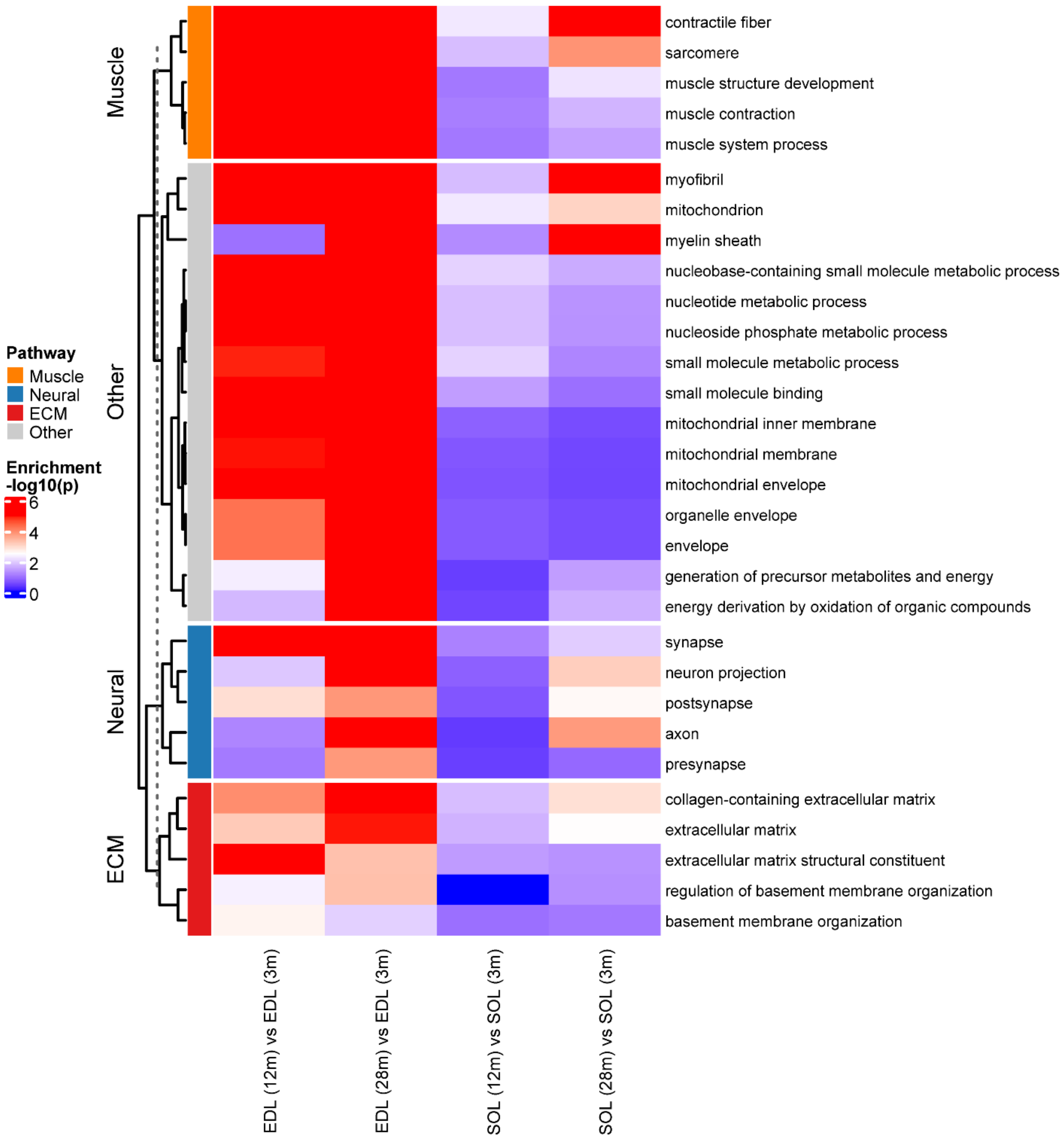


**FIGURE S3**. Heat map of Gene Ontology (GO) term enrichment across aging-related muscle comparisons. Each column represents a differential gene set derived from a pairwise comparison between time points in SOL and EDL for decoupled genes. The top 30 most enriched GO terms (based on adjusted p-values) were selected, with a minimum of 5 terms per major functional category (Muscle, Neural, ECM). Colors indicate the significance of GO enrichment (-log10 p-adjusted), with blue denoting higher enrichment in the earlier time point of each comparison and red indicating higher enrichment in the later time point. Muscle-related processes are predominantly enriched in early SOL transitions and decline with age, while neural and mitochondrial terms show late-stage enrichment in EDL, reflecting divergent aging pathways and tissue-specific vulnerability.
